# Brain Network Constraints and Recurrent Neural Networks reproduce unique Trajectories and State Transitions seen over the span of minutes in resting state fMRI

**DOI:** 10.1101/798520

**Authors:** Amrit Kashyap, Shella Keilholz

## Abstract

Large scale patterns of spontaneous whole brain activity seen in resting state functional Magnetic Resonance Imaging (rsfMRI), are in part believed to arise from neural populations interacting through the structural fiber network [18]. Generative models that simulate this network activity, called Brain Network Models (BNM), are able to reproduce global averaged properties of empirical rsfMRI activity such as functional connectivity (FC) [7, 27]. However, they perform poorly in reproducing unique trajectories and state transitions that are observed over the span of minutes in whole brain data [20]. At very short timescales between measurements, it is not known how much of the variance these BNM can explain because they are not currently synchronized with the measured rsfMRI. We demonstrate that by solving for the initial conditions of BNM from an observed data point using Recurrent Neural Networks (RNN) and integrating it to predict the next time step, the trained network can explain large amounts of variance for the 5 subsequent time points of unseen future trajectory. The RNN and BNM combined system essentially models the network component of rsfMRI, and where future activity is solely based on previous neural activity propagated through the structural network. Longer instantiations of this generative model simulated over the span of minutes can reproduce average FC and the 1/f power spectrum from 0.01 to 0.3 Hz seen in fMRI. Simulated data also contain interesting resting state dynamics, such as unique repeating trajectories, called QPPs [22] that are highly correlated to the empirical trajectory which spans over 20 seconds. Moreover, it exhibits complex states and transitions as seen using k-Means analysis on windowed FC matrices [1]. This suggests that by combining BNMs with RNN to accurately predict future resting state activity at short timescales, it is learning the manifold of the network dynamics, allowing it to simulate complex resting state trajectories at longer time scales. We believe that our technique will be useful in understanding the large-scale functional organization of the brain and how different BNMs recapitulate different aspects of the system dynamics.

## Introduction

Over the past decade, our understanding of spontaneous whole brain activity and coordination between brain regions has largely been obtained through non invasive resting state functional Magnetic Resonance Imaging (rsfMRI) studies [23, 31, 37]. Resting state, a state without an explicit task or stimulus, has surprisingly complex whole brain trajectories that are well structured and highly dependent on the previous brain state [1, 5, 29, 37]. Current generative models such as Brain Network Models (BNM) attempt to characterize whole brain activity as the interaction between a single neural population and the activity of its network neighbors defined by its structural fiber connections as measured through diffusion tensor imaging (DTI) [7, 18]. Although there are many variants of the model that use different sets of differential equations to describe the activity at each node, all Brain Network Models heavily rely on the description of the structural network through which they interact [27].

Many types of Brain Network Models are able reproduce time averaged properties of rsfMRI such as average functional connectivity (FC) that is defined as the correlation between brain regions over long periods of time. The time averaged properties are thought to be directly related to the structural network [30]. However, they are worse and more variable at reproducing transient dynamic features that occur at shorter timescales and depend on the exact description of the differential equations [8, 20]. We propose to synchronize BNMs to empirical data by solving for the initial conditions of the model using modern machine learning techniques. This approach will allow us to better distinguish between models based on their ability to reproduce specific dynamics observed in the data, since they are all able to produce average measures such as functional connectivity [8].

In this manuscript, we demonstrate a method using recurrent neural networks (RNNs) to learn the initial state of the Brain Network Model and then predict future moment to moment changes in the rsfMRI signal. RNNs with constraints from biology have been recently shown as an efficient tool in solving for and modeling unknown systems of differential equations [9, 25]. By applying this technique, we can quantify how much of the variance of future resting state activity can be accounted for through nonlinear network-based propagation as opposed to other sources of activity such as from a stimulus.

For our dataset, we used fMRI scans from 447 Human Connectome subjects [34] using both task and rest to train a sequential Autoencoder, a model that is trained to predict one time step into the future. Unlike a regular Autoencoder, the latent state in our model is represented by the variables in the Brain Network Model. We implement two variants of this model that have different latent states, the Firing Rate and the Wilson Cowan model [7, 27, 36]. After training, the system was tested on a set of 40 unrelated unseen subjects to see how well the single step forecasting generalizes to the population. We then extend our model to determine how many more timesteps we can predict accurately into the future. We also examine the errors across each node of the network to see if structure exists in the distribution of errors across different regions of interest (ROIs). Finally we test long periods of simulations of our generative model in a similar manner currently used for evaluating traditional BNM, in order to see whether it can reproduce dynamic properties that occur over minutes.

This Brain Network Autoencoder (BNA) method offers three main strengths in comparison to other methods that are currently used to simulate whole brain signals:

1. It solves the problem of comparing simulated and empirical data without using time averaged metrics such as average FC, by directly using real data to initialize the model and by measuring differences in predicted the transient dynamics on a moment to moment basis.
2. It allows us to estimate latent variables such as firing rate or excitatory and inhibitory currents that can be verified using multimodal recordings.
3. In long simulations of the BNA, the simulated signal exhibits dynamic properties seen in empirical rsfMRI that occur over the timescale of minutes, that are not reproducible using traditional BNM techniques.

Therefore, we believe that the Brain Network Autoencoder (BNA) will be a useful tool to help us understand brain dynamics at the macroscale level.

## Results

### One Time Step prediction

For this application the BNA is trained to predict one time step in advance. In Figure 1, we present the results of predicting the next time-step from the previous time-step for the two different variants of BNA, the Firing Rate Model and the Wilson-Cowan model. Although both are able to reproduce the spatial-temporal signal as shown in Figure 1 (middle top) and Figure 1 (middle bottom), they differ in the latent or hidden variables used to represent the transitions. For the Firing Rate BNA, the measured data is projected into a space with firing rate as the hidden variable for each region, as can be seen in Figure 1 (right top). The latent variable time series has a high degree of similarity to the original signal (correlation *>*0.9) shown in Figure 1 top right. The latent state is then passed onto the BNM which integrates it according to the Firing Rate model to predict the next time-step. The traces of the input, output and latent state for a single ROI is shown in Figure 1 (left top). For the Wilson-Cowan model Figure 1 (bottom row), the latent state is represented by two variables, excitatory and the inhibitory currents, and their interaction through the Wilson-Cowan model produces the next rsfMRI time step. The excitatory current is positively correlated with the measured signal and the inhibitory current is negatively correlated with the signal although both to lesser degrees than the Firing Rate model. The models perform relatively similarly in predicting one time step in front and are able to reproduce the input signal with an r-squared of 0.95 averaged across all areas.

**Figure 1.**
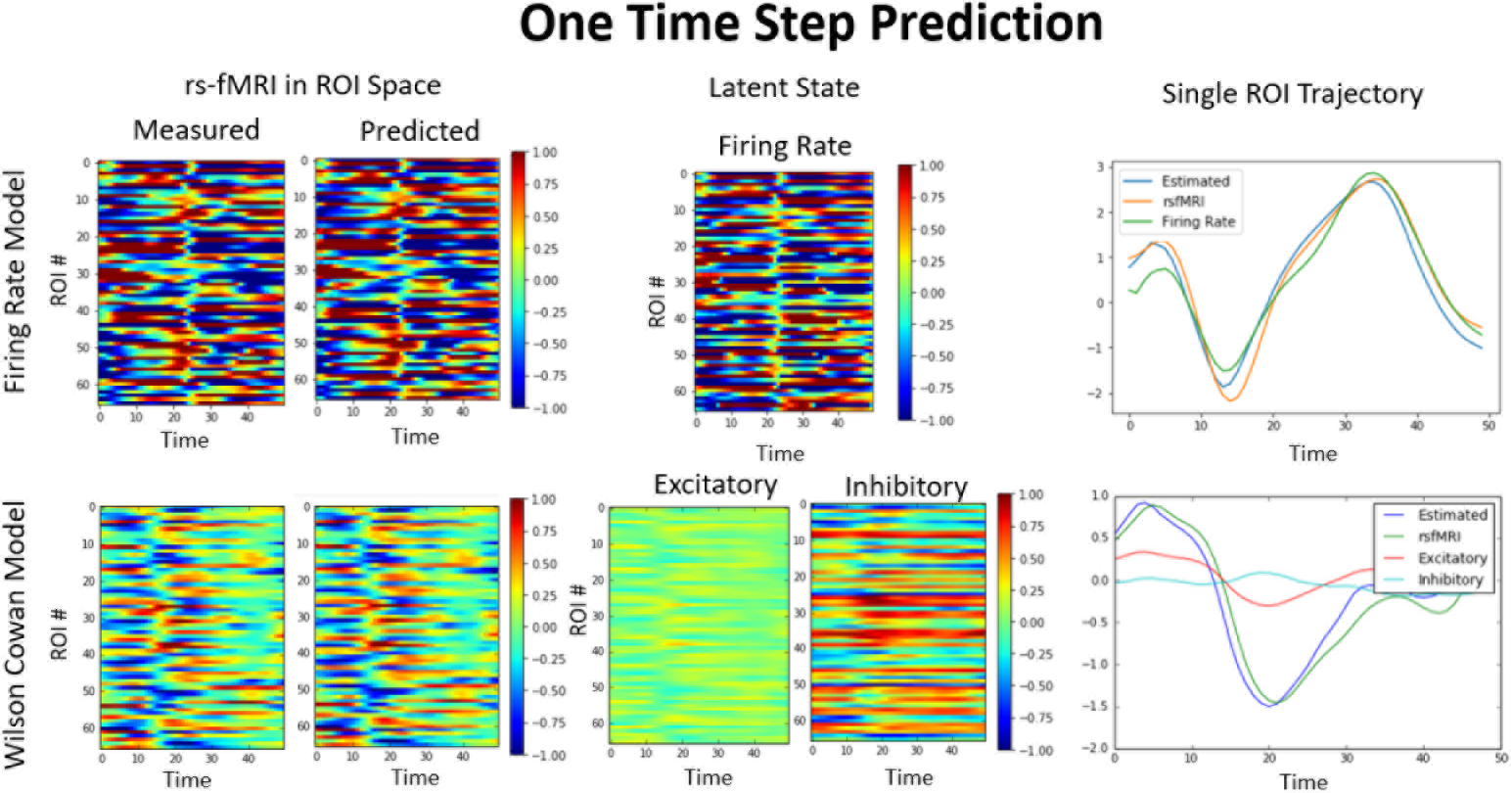
One Step Prediction. Two different Brain Network Autoencoders used for a single time point prediction. The Autoencoder takes as input the measured signal (left most) at time step t and outputs the predicted (second from Left) signal at t+1. The Autoencoder projects the input into a space constrained by the Brain Network Model Equations (middle panel) using either as state variables in the Firing Rate model or the Wilson Cowan model, which are then integrated to produce the predicted output. The right most panel shows the timeseries of a single ROI for the input, output and latent state. At one time step the accuracy in terms of r-squared across all ROIs is on average 0.95. The time axis is in unit TR, which is 0.72 sec.

### Multiple Time Step prediction

The sequential Autoencoder can also predict multiple steps into the future by recursively feeding the predicted output in as the next input. The performance of multi step time forecasting is shown in Figure 2 (top left), where the averaged r-squared across a test and a subset of the training data of the same size for both BNA variants are compared with a naive variant of the Autoregressive model (ARM) that assumes the next time point is the previous time point (see methods). The ARM is similar to the current approach used to differentiate task from rest signals, where the generalized linear model takes the time steps before task activation and uses them as a regressor to remove the resting state activity from task responses [31]. Although the ARM performs as well as the BNA for the first time point, the BNA is able to reproduce the first three time steps with an r-squared of around 0.9 or higher as opposed to ARM model which is only greater than 0.9 for the first time step. The test and training performance is relatively similar for the Autoencoders, only when all the parameters are set correctly and the network is not over or under trained (see Methods Figure 6 for more detail).

**Figure 2.**
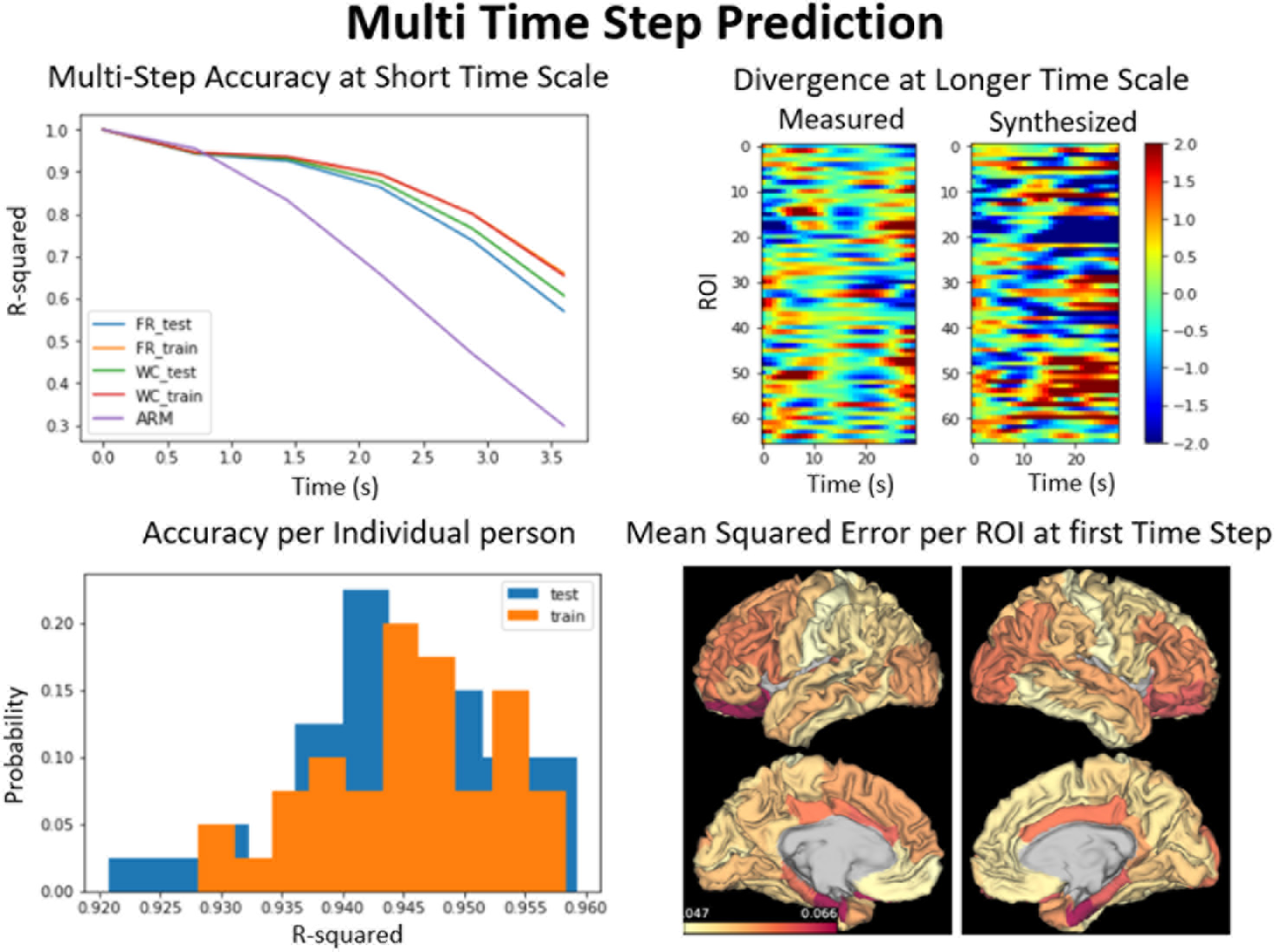
Error across multi-step prediction. Top Left: Accuracy of our generative model in synthesizing the first few time points. The accuracy of the Firing Rate and Wilson Cowan model are compared on training and test data sets and to the autoregressive model (ARM). The error compounds and gradually increases until the model diverges completely from the measured signal around 10 seconds and continues along its own dynamics (top right). Bottom left: Histogram of r-squared for each individual in test and train data sets shows that it generalizes across individuals. Bottom right: The mean squared error (MSE) for each region of interest (ROI) in predicting the first time step. The MSE is used here to compare differences across ROIs, because it was the error that was used to train the system and is more reproducible across instantiations.

Characteristic of Autoencoders, the error compounds at every timestep, because the previous errors are propagated to the next time step. This causes the model to completely diverge by 10 seconds from the measured signal as shown in Figure 2 top right. The bottom left figure also shows that the BNA generalizes across individuals, as the histogram of the errors is roughly the same for all the individuals in the training or the testing data set. The two different BNA variants, the Firing Rate and the Wilson Cowan are similar in performance as seen in top right of Figure 2, with the Wilson Cowan having on average a higher r-squared on the test data set. The BNA does not perform equally in predicting each of the ROI timeseries. It predicts certain regions with a higher accuracy than the others. Th mean squared error per each ROI for the first time step is shown in Figure 2 bottom right. The Mean squared error was used here instead of r-squared, because the network was trained to minimize this gradient during training and most accurately represents the performance on each ROI. The error was largest in the ROIs in the temporal lobe, namely the entorhinal cortex, parahippocampal gyrus and the temporal pole. These regions are the least connected to the rest of the network and more connected with subcortical regions that are not simulated [7].

### Analysis of Long Simulations

In order to assess properties of the simulated signal at longer periods, the BNA was used to generate 1000 time points or 12 minutes of data. Properties of longer simulations of BNA were compared to those of the empirical signal. In Figure 3, the average Functional Connectivity and the Power spectrum of the empirical and the BNA as well as a traditional Firing Rate BNM is compared. The traditional BNM FC has a weak correlation with the empirical FC (0.35) and is in the range of most traditional methods (0.3-0.6) [8, 20, 27, 28]. The BNA performs much better at reproducing the detailed relationship between ROIs seen in FC, compared to the traditional model where groups of ROI are synchronized over long periods of time causing a blocky patches in the FC when the ROIs are ordered by highly connected subgraphs [7, 20]. The FC of the BNA has a high correlation of 0.83 (Firing Rate) and 0.7 (Wilson Cowan) to the actual measured signal. The spectral power of the empirical and the simulated signal are indistinguishable in the range 0.01 to the 0.3 Hz, and has the characteristic 1/f linear slope of around 0.9. The traditional BNM which has less temporal structure and broad band power in the lower frequency and then falls off more sharply after 0.1 Hz [20]. At higher frequencies the model tends to produce much higher levels of noise than the empirical signal and the traditional BNM both which have already been filtered in the pre-processing steps. Before analyzing the simulated signal with dynamical analysis techniques, we therefore filtered it at 0.3 Hz to minimize the high frequency power that would interfere with the dynamic analysis algorithms.

**Figure 3.**
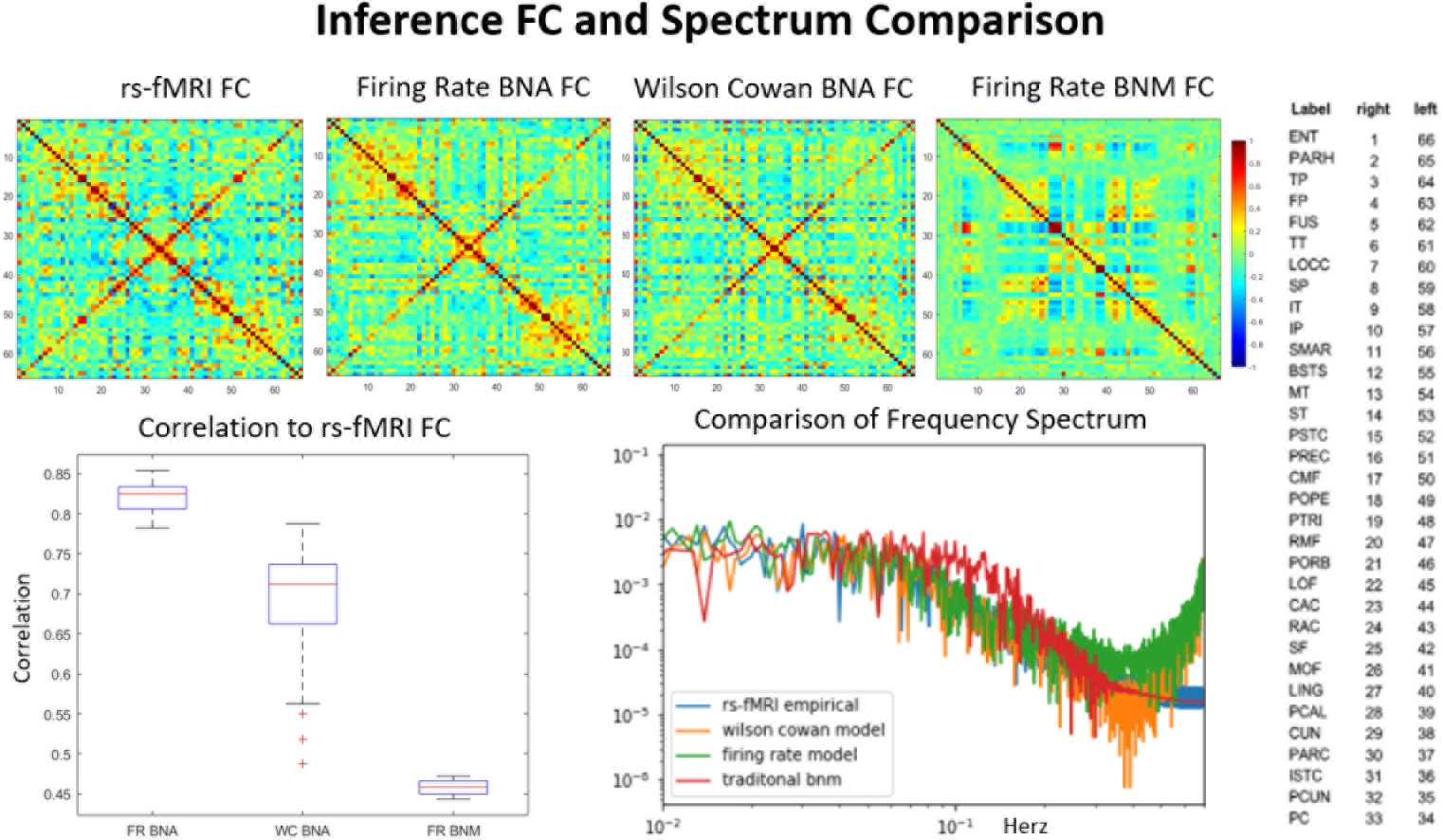
Average Functional Connectivity and Power Spectrum. Comparison of average functional connectivity from empirical rsfMRI (top left), BNA Wilson Cowan (top middle), BNA Firing Rate (top right), and a traditional Firing Rate BNM (bottom left). The simulated FC matrices have a high degree of correlation 0.7-0.85 with the empirical FC unlike the traditional BNM which have a correlation of 0.5. Each axis in the FC plots represent the regions in the ROI which are shown on the right. The frequency spectrum (bottom right) of the BNA follows that of the empirical signal exactly except at the higher frequencies (*>*0.3 Hz) where the simulated signal has much larger power. The traditional BNM has less structure in the frequency range (0.01 - 0.1Hz) and has equal power in most of the range compared to the rsfMRI and the BNA models. The traditional BNM and the empirical signal also have been filtered at 0.3 Hz and while the BNA models are not.

We also analyzed the simulated signal for unique trajectories known as quasi periodic patterns (QPP), which could also be considered a limit cycle [22]. Limit cycles are a property unique to non-linear systems, and reproducing such a property would mean that the generative model reproduces some of the dynamics features of rsfMRI despite its divergence from measured signals. In Figure 4 we have plotted the QPP pattern for the rsfMRI signal (top left), a traditional generative Firing Rate BNM model (bottom left) and both the BNA variants (top middle and right). The empirical QPP pattern involves a twenty second trajectory that switches from task positive networks (first half of the template) to the more internal or default mode networks of the brain (second half of the template) [22]. After phase adjusting the templates, the maximum correlation of the Firing Rate BNA QPP was 0.75 and the Wilson Cowan BNA QPP was 0.43 to the original template. This is very different than the traditional dynamics seen in BNM (bottom left) which produce blocky limit cycles, of clusters of nodes that are highly synchronized together and activating together. The BNA produces QPP that are highly structured spatially and temporally. The correlation between the QPP template and the signal is plotted in the bottom middle, where certain time points show high degree of correlation to the trajectory in the QPP template. Thresholding at 95 percent significance, the occurrence of these QPP patterns is around 1.3 times a minute in the rsfMRI data. The BNA models have similar rates, where the Firing Rate BNA has a occurrence of 1.19 times a minute. The Firing Rate BNM model shows more variance in the number of QPP cycles per minute (bottom middle Figure 4).

**Figure 4.**
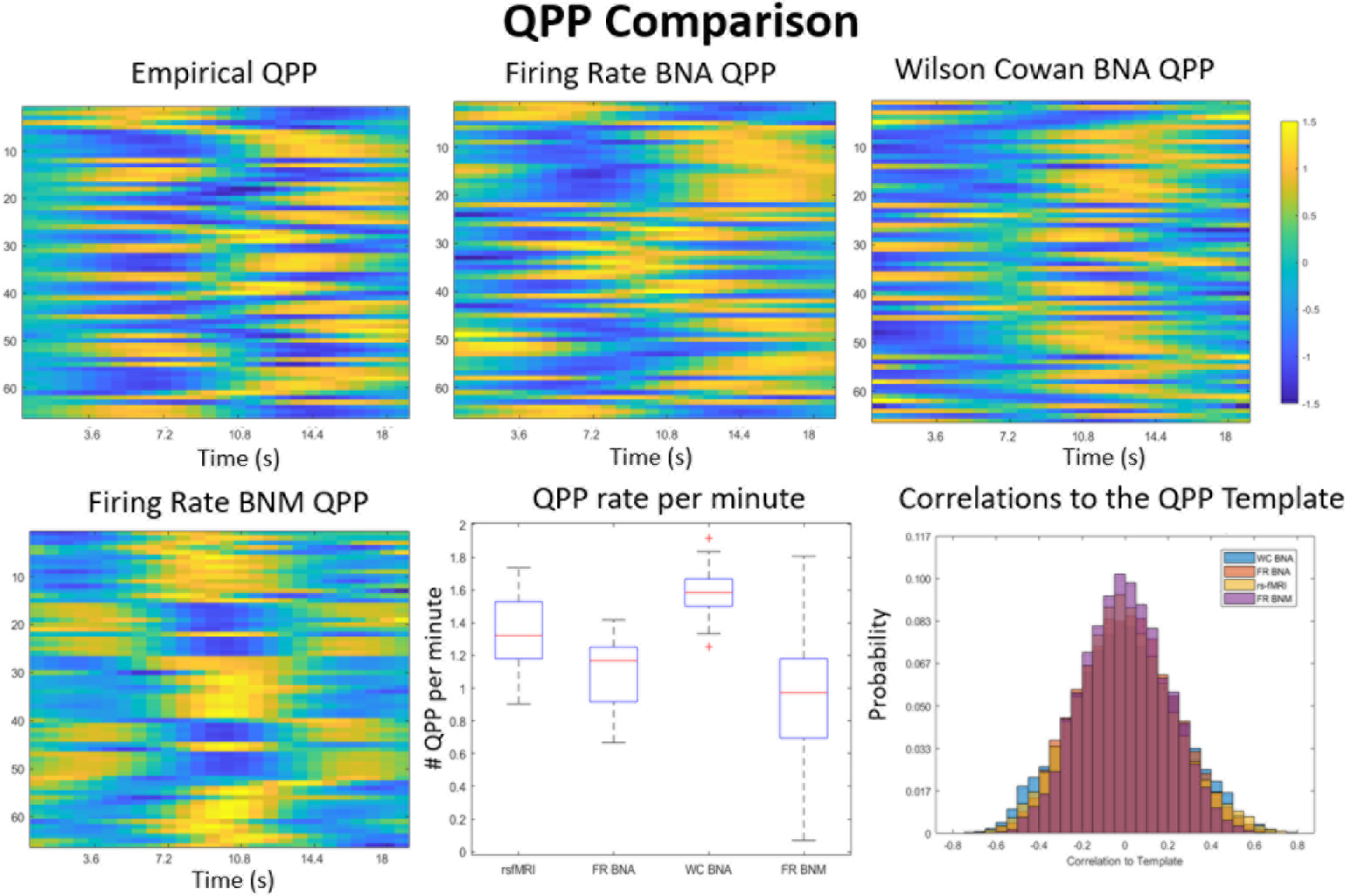
QPP Template Comparison. Comparison of the different QPP Templates is shown in the top row between measured data (top left), the Brain Network Autoencoders (BNA) (top middle and right), and the older BNM (bottom left). The QPP template represent a unique 18 sec trajectory of all the ROIs (y axis) that repeats itself on average 1.3 times per minute (bottom middle). The rsfMRI signal is highly correlated with the template for during specific time-points in its trajectory as seen in the distribution of correlations to the template (bottom right). The Wilson Cowan and the Firing Rate BNA have similar distributions, while the BNM template is least correlated with its own data. The Firing Rate BNA QPP is the closest to the empirical QPP (correlation 0.73) and occurs roughly 1.19 times per minute. The Wilison Cowan BNA QPP occurs a little faster around 1.4 times a minute and has a correlation of 0.43 with the original template. The older BNM QPP is more of an on-off trajectory and does not have the intricate delays and temporal structure as seen in the QPP of the empirical signal or the BNA models.

Another property of rsfMRI that has been studied is the existence of brain states, which can be described as large scale patterns of functional organization that are stable over the span on the order of around 40 seconds [1, 21].The brain transitions through these states over time [1]. Algorithms such as k-Means have typically identified 6-7 states. We applied k-Means clustering on short windowed functional connectivity matrices (50 sec) to find these states in the simulated data (see methods for more detail). In Figure 5, we show the comparison between our BNA models, the Firing Rate BNM models and the measured signal for cluster centers as a result of the k-means algorithm. We quantified how close the centers are to each other, by taking the maximum correlation of each center to those measured in rsfMRI. We calculated the length of time in each state (top left), the transition likelihood between states (bottom middle) and how many unique states were observed in a single scan (bottom left). The centers of the BNA models (middle two) compared to the traditional BNM (right most) are much more distinct from each other. The Firing Rate BNA model has the highest correlation with the rsfMRI states (0.8 on average) and a similar number of states seen during a single scan. It does has have much lower dwell time and seems to move between states twice as fast as the measured signal. The Wilson Cowan has more variable and diverse centers and tends to have fewer of them in a single scan, but tends to dwell in them around as long as the measured data. The traditional Firing Rate BNM model is the least accurate, has few transitions between states, and dwells in a single state for a very long time.

**Figure 5.**
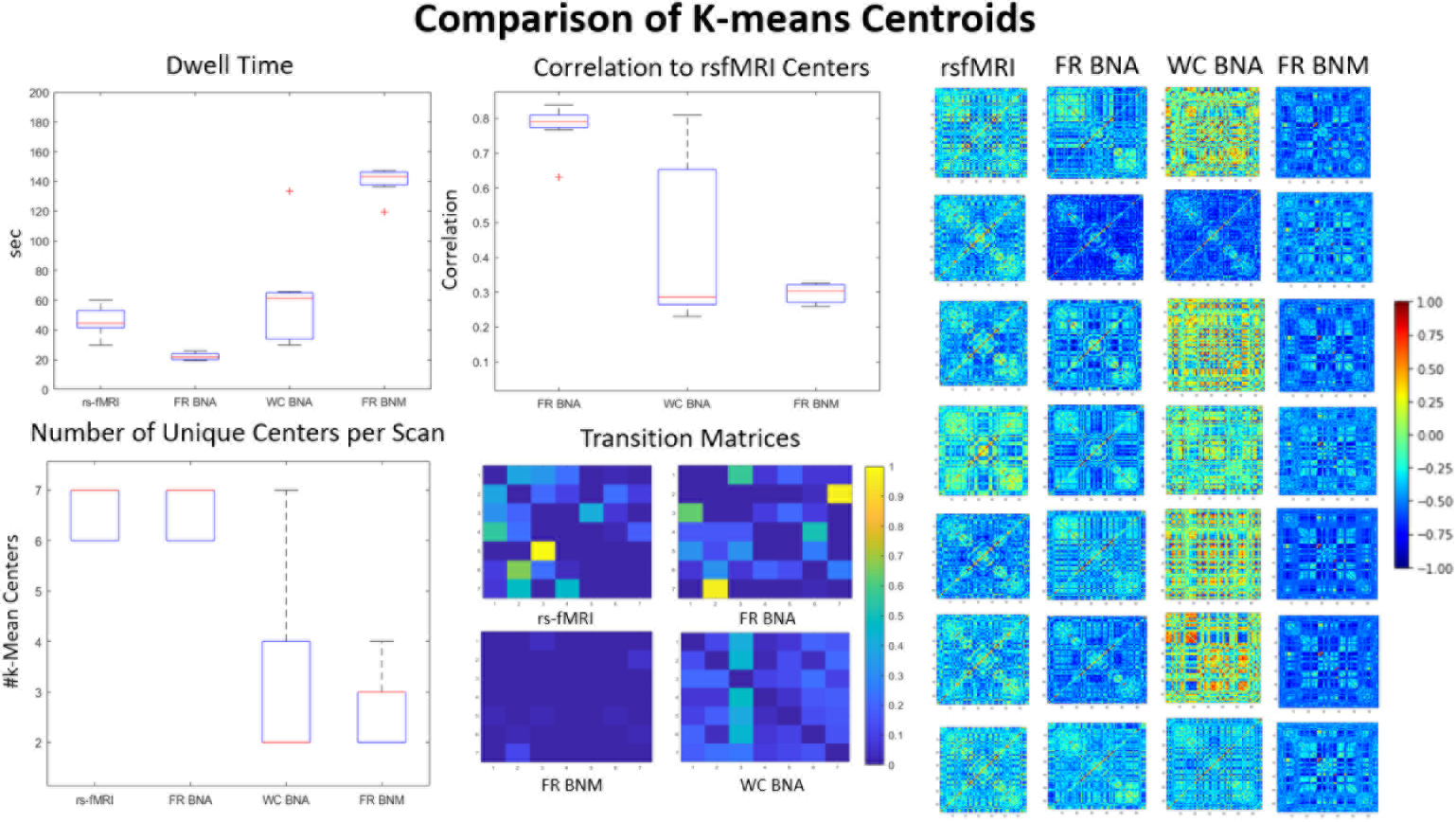
k-Means Comparison. This figure compares the k-means centers and the transitions for the simulated (BNA and naive BNM) and the empirical signal (30 scans of 15min). The seven centers are shown in the far right for each category (FR: Firing Rate, WC: Wilson Cowan). The a boxplot of the max correlation (middle top) of each of simulated centers to the centers from the rsfMRI data. The dwell time in seconds in each of these centers is shown top left. The rest of the transition probabilities (diagonal zeroed out) are shown bottom middle. The number of centers in each of the 30 scans is also variable even though they all are defined to have seven clusters across all scans (bottom left).

## Discusion

In this manuscript, we adapted the Brain Network Model with the RNNs in order to make short time future predictions from observed rsfMRI. Using this approach, we showed that much of the moment to moment variations can be explained through network based propagation of the previous measured rsfMRI data point. We then showed that based on learning the initial conditions from past rsfMRI data, the system can generate resting state trajectories that recapitulate over larger time scales.

### Predicting Moment to Moment Variations

We showed that a network-based model can account for up to 95 percent of the variance in the fMRI signal between two adjacent time points. This reproduction is not unique however, and can be estimated using any number of latent variables. Although more complex architectures such as Variational Autoencoder might be able to successfully predict future rsfMRI data [25], the BNM provides an adequate rough guess of the system dynamics for the Autoencoder to converge. This is most likely because the gradient vector evaluated at every time step is relatively close to the structural matrix because the structural matrix represents the physical avenue to propagate information between regions. This information helps the model to converge during training and make accurate predictions. In fact, our simulation of a RNN with no Brain Network Model failed at minimizing the error between the predicted and the measured data point. Moreover, unlike a traditional Machine learning approach, this approach yields testable latent variables that can be further evaluated using multimodal datasets, i.e Magnetoencephalography (MEG) recordings that have been used to generate excitatory and inhibitory currents synchronized with concurrent rsfMRI recordings [26].

Fluctuations in spontaneous whole brain activity have been shown to be non-random and highly structured [37]. This suggests rsfMRI has both deterministic and stochastic components. The variance explained by the BNA at one time prediction represents a lower bound of the amount of determinism that exists in the signal. It is not surprising that this is the major component of rsfMRI since the signal has shown to be highly auto correlated with itself [2]. The simplified Autoregressive model, which assumes a steady baseline at the last measured timestep, has similar results in performance to the BNA when compared to a single timestep and has an r-squared of 0.97. However, for multiple timesteps into the future the Autoregressive performs poorly, compared to the BNA models. The two different BNA models perform at short term scales about as well as each other. This suggests that the trajectory in the short time span is predictable to a certain degree regardless of the approach, but thereafter it starts diverging from the empirical measurements. The divergence from the original trajectory could be due to a number of sources, such as unknown task or stimulus information, noise, or simply a mismatch between the algorithm and the data that increases over time. Note the BNA itself is not a deterministic system. The latent space variables are modeled as distributions before they are sampled resulting in a stochastic system. However, our results show that network based activity is the dominant component of dynamics at short time scales and is mostly deterministic and learnable.

### Evaluation on Long Term Dynamics

Although both rsfMRI and the BNA models are stochastic, long term simulations of the network based model are able to reproduce trajectories that are similar to those seen in rsfMRI. Individual trajectories are varied but they repeat over time, suggesting that rsfMRI follows a bounded stable manifold which the model is able to estimate.

Therefore random walks across this manifold have shared properties in both the model and the empirical signal. Our results also suggest that most of the resting state manifold is strongly related to the network-based activity rather than input or random perturbations from noise sources such as higher neural processing.

The strongest metric demonstrating this relationship is average FC which has a large correlation to the empirical dynamics (0.9 *>*correlation *>*0.8). This is unsurprising since the traditional BNMs do almost as well as the BNAs in this metric, and correlations as high as 0.7 have been reported in literature [28]. Average FC seems to be more related to the structural input than the description of the dynamical system [8, 20]. However, the BNA does better than most BNMs in estimating interhemispheric FC correctly, which is usually challenging in network based models because there are far fewer interhemispheric than intrahemispheric connections detected with diffusion MRI. The power spectrum profile is also mostly reproducible by the model, except in the very high frequency where the model has a lot more power than the empirical signal. This might occur due to the lack of friction in our model, namely that the signals are constantly propagated through feedback loops in the network without loss of energy, unlike the real system. Since most of predictability of resting state comes from the structured low frequency activity, we can filter synthesized signal without losing too much information. Other traditional BNM using the virtual brain have also reported similar performance on power spectrum profiles [26].

Although most traditional BNM have been able to reproduce to some degree the long-term averaged properties such as average FC and power spectrum, they have had a harder time in reproducing faster scale dynamics such as reoccurring unique trajectories or the multi-state transitions seen in dynamic FC [8, 16, 20]. The results from the QPP analysis, which extracts limit cycles, show that the simulated signal has a similar 20 sec trajectory and that pattern is repeated over the course of minutes. The results from the k-Means analysis on time varying FC matrices, shows that the simulated signal has similar state transition in terms of both number and the spatial patterns to those seen in empirical rsfMRI. This suggests that both of these properties arise naturally in the correct non-linear network-based representation of rsfMRI which can be inferred from the data using machine learning techniques. The Firing Rate BNA seems to fit the data better than the Wilson Cowan BNA. This might be due that the Wilson Cowan BNA has additional non-linearities due to the interaction between the excitatory and inhibitory currents.

A direct comparison between our model and other BNM models in literature on complex dynamical metrics is difficult because most BNM models use their own unique metric to compare against rsfMRI and there is no established standard. The origin of these complex dynamics have been explained in different theoretical ways. These complex transitions can arise to the particular non-linearities of the system [16], which can result in multiple attractors and limit cycles naturally. They can result from parameter changes to the network strength or Hopf bifurcations that cause the system to change its dynamics over time [10, 28]. They can also be the result of adding external input and stimuli into the system causing a change from the zero-input manifold and altering the dynamics [3, 11]. These are not mutually exclusive and could induce the changes at once. Our implementation, is closest to the first interpretation of rsfMRI. We explain the observed non-linear properties of the data purely based on network propagation without the need of external input or a change of a bifurcation variable.

### Errors across different ROIs

The error in predicting dynamics is not evenly distributed across all regions of interest. The error in reproducing the dynamics at one time step is highest in the nodes of the Limbic system (Figure 2 bottom). We believe that our model performs less accurately in this system because they are highly connected to the amygdala and the hippocampus, which are not simulated in the model, and are the least connected nodes to the rest of the network [7]. Moreover, tractography has also been known to underestimate the Uncinate Fasciculus, the major highway between the temporal lobe and the frontal areas, which forms the backbone of the limbic system. The fiber has a very sharp angle which is hard to follow using tractography [33]. The echo planer imaging (EPI) sequence used to obtain rsfMRI data, has also known susceptibility issues at interfaces, which would affect the nodes at the proximity such as the frontal pole and the temporal pole, both which have larger mean squared error compared to the other nodes.

### Comparison to other Machine Learning and Time Forecasting models

Similar time forecasting has been attempted or is being attempted by several different labs at a the time of this manuscript. A variant utilizes a Variational Autoencoder to find a latent space of brain trajectories that would fit the current data [6]. Another RNN-ICA version uses ICA vectors as the latent space, while another method uses Hidden Markov Model to model the hidden states [17, 35]. However, our method is unique in using Brain Network Model as a latent space, whose variables are more interpretable since they represent the state of each neural populations activity and can be tested using multi modal data. Moreover, none of the other architectures use their model for time series foresting or dynamical analysis, hence their results are not directly comparable to our work, although their methods are similar.

### Limitations

There are many assumptions that limit the scope of our approach. Machine Learning although good at learning structures in data sets, has a shortcoming of arbitrarily creating a system to fit the data and every instantiation of the system produces slightly different properties of the simulated system. We tried to address this, by using various techniques such as using structural constraints, dropout of Long Short Term Memory (LSTM) units, using probability to track the latent variables, and taking the results of multiple runs, in order to make the system more reliable and reproducible. Another limitation of this model is that it needs 50 time points prior to the data point, in order to solve for the initial conditions. Shorter time intervals than 50 time points are faster to train, but are less accurate in estimating slow processes. The longer segments required a larger LSTM network and longer training times and were less accurate in our dataset. There are more complex architectures that could solve for the initial conditions faster, such as a forward-backward LSTM architecture [25]. On the network side, the parcellation scheme reduces the complexity of the signal and discretizes the network. Improvements can be made by allowing for continuous propagation along the cortical sheet, as in the neural field models. Tractography also has its limitations, and better estimates of structural networks should make the model more realistic and improve results especially in regions that are not very strongly connected to the rest of the network. Simulating more of the central nervous system including sub-cortical regions would also lead to a more biologically plausible model.

### Conclusion

We set out to investigate the extent to which network based theory can explain the moment to moment variations seen in rsfMRI signal. Using a novel Machine Learning approach, we solve for the initial state of traditional network based models, and show that we can account for most of the variation seen in the signal and predict accurately (*>*0.6 r-squared) for at least 5 consecutive timepoints. Longer instantiations of the system shows that our model is able to produce complex trajectories of the non-linear dynamical system on the order of minutes. We believe that our BNA will be useful when a generative model of rest is needed. Moreover, it can be trained to predict in real time, which allows contrast against dynamics that contain deviations from rest such as in task fMRI studies. In the future, it can also be used to investigate deviations from the manifold such as in task input or due to noisy sources.

## Methods

### Mathematical Background

The Brain Network Autoencoder is constructed using the constraints from the Brain Network model, in conjunction with a recurrent neural network variant known as Long Short Term Memory (LSTM). The overall design is shown in Figure 6 and implemented using Python Tensor Flow. The architecture is a sequential Autoencoder, as it is trained with the previous time point to predict the next consecutive time point and uses a latent space where the dynamics are constrained to a smaller space defined by BNM equations to reconstruct the next time point.

**Figure 6.**
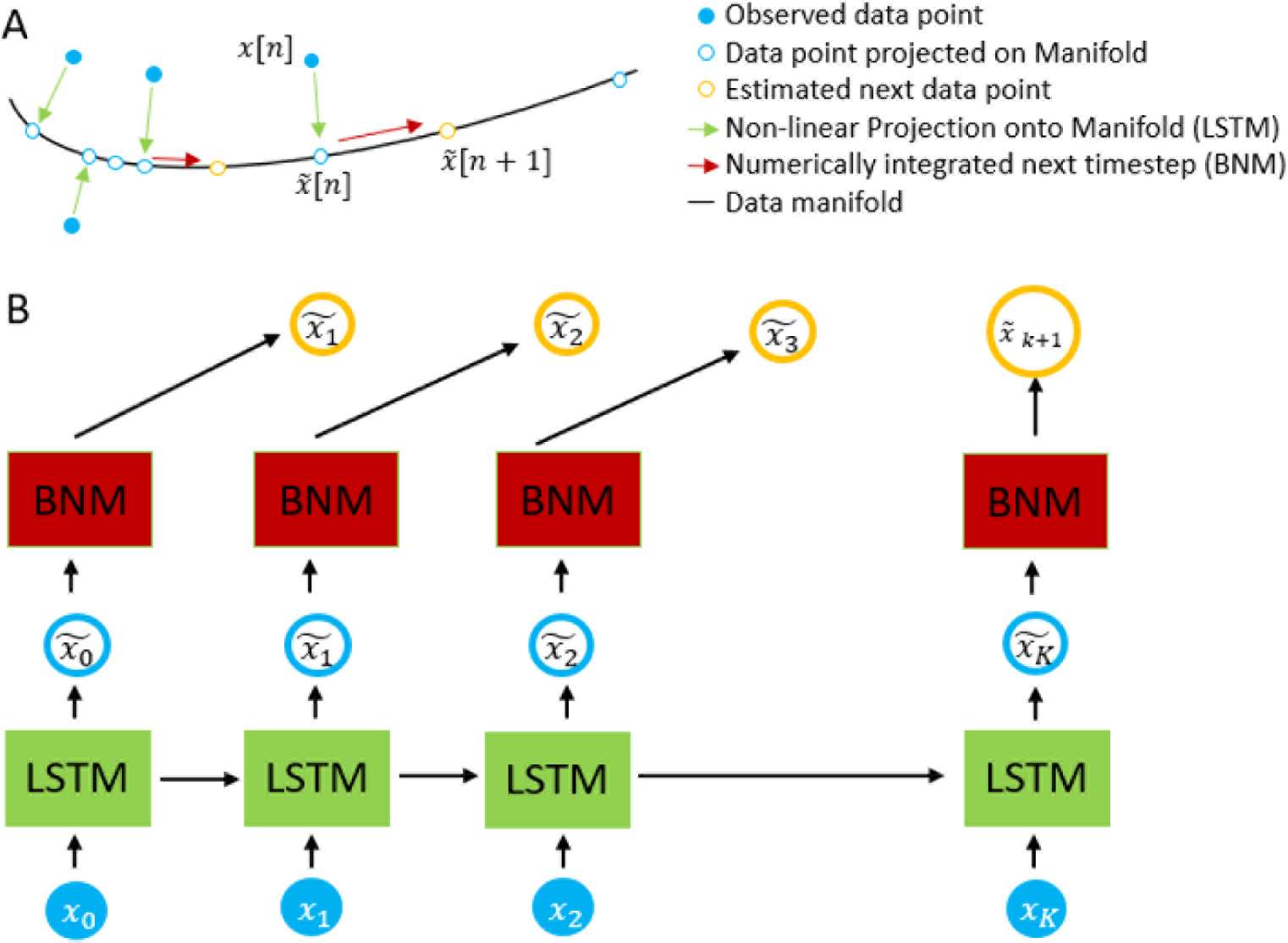
Schematic of the Autoencoder. The measurement *x*(*n*) is passed into the LSTM in order to estimate *x*(*n*) which lies in the data manifold. Using the BNM forward equations and *x*(*n*) as our initial conditions, we estimate *x*(*n* + 1). The system is trained by difference in our predicted vs actual measurement at *x*(*n* + 1).

Formally, in order to predict the next time point, for each neural measured time point x(n) we map it to the space *F* (*x*(*n*)). F is the transformation performed by the RNN and lives in *R^M×M×T^*, where M represents the M distinct ROIs being modeled and T is the length of previous time-points the LSTM depends on. The next time point is computed as *x*(*n* + 1) = *BNM* (*F* (*x*(*n*))). In essence, the LSTM does a non-linear coordinate transform of the vector *x*(*n*) into the Brain Network Space where the dynamics are well defined and we can predict the next time point. This process shown pictorially in Figure 6a, where we show the projection of each data point shown in filled blue circle into the manifold represented by the BNM shown in a hollow blue circle. On the manifold, we can use the BNM equations to update it to the next time step shown in orange. Fig 6B shows the actual architecture used to update the timesteps.

For the simplest implementation of BNM, the firing rate model, we can assume the function to be a linear with the observation such that BNM(x) becomes *A x* where the matrix *A* is the graph Laplacian and *A* = *k SN I*, where *k>*1 and SN is the structural matrix as measured through tracktography using diffusion tensor imaging (see methods section structural matrix) [15]. We use graph Laplacian, because they represent a well studied dynamical system known as the consensus equation. On its own, the consensus system does not add in any unstable dynamics due to all of its eigenvalues being less than zero, if k is set to less than 1 [24]. Therefore the network propagation dies out over subsequent timesteps. The eigenvectors of A have also shown similarities to rsfMRI networks [4]. This algorithm assumes that the Jacobian matrix representing the changes of one brain region with respect to another more or less lies in the direction of the structural fiber network and the non-linear discrepancies are dealt by the LSTM [18]. This can be seen in the results section (Figure 1) where latent space of the Firing rate model is almost identical to the measured data, suggesting the transformation is near an identity transformation. In a more complex BNM, such as the Wilson Cowan, the excitatory current is strongly correlated with the signal although less than in the Firing Rate model, as the model has its own inbuilt non-linearities and deviates further from the graph Laplacian.

### Implementation

The preprocessed data (see methods section Preprocessing) is first cut into contiguous segments of length k. This whole segment is then passed into the Long Short Term Memory unit as shown in Fig 6 B. The units are built using the Tensor Flow API, specifically the GPU boosted version to improve speed and performance. The LSTM units take in a series of consecutive time points, and output a sequence of the same length of time points. The LSTM units are a form of recurrent neural networks and have memory of previous time points by using a hidden state vector which it uses as a input to itself for the next consecutive time point. Hence, LSTMs have become popular in the Machine Learning community because of their success in using this architecture in modeling time series such as speech and natural language processing, in self driving cars, and even in neural Turing computers thought to emulate biological intelligence [12–14]. Moreover, they solve the problem of learning structure across infinite sequences of consecutive time points by using a forget gate to truncate inputs seen from a long time ago. In practice this means that they need to be trained with a finite sequence length of data.

For our implementation we tested data of length 25, 50 and 100 time points (18, 36, 72 sec) as seen in Figure 7 left. The model performed best on 50 length segments, and slightly worse for shorter and longer segments. The LSTM network was also stacked into several layers in a similar manner that convolutional neural networks are stacked together in a series. We used 7 identical layers to model the fMRI timeseries. In general, more layers improve accuracy as long as there is enough data in the training set to scale the size of the network, otherwise there is a risk on overfitting. Using the inference error as a metric we also swept the number of training iterations until the performance on unseen crossvalidated testing data was about the same as the training data as shown in Figure 7 right. For the crossvalidation we split the data of 447 individual scans for 40 test and 407 training samples randomly. At the right amount of training steps, the system does relatively equally in test and training sets. An overtrained or undertrained network on the other hand, resulted in large differences in test and training, although all three models do equally well on the training dataset. In order to additionally control for overfitting, we also used the inbuilt tensor flow dropout function that prunes a large number of the weaker weights used in the LSTMs. This has been shown in neural networks to better generalize to unseen test data [32].

**Figure 7.**
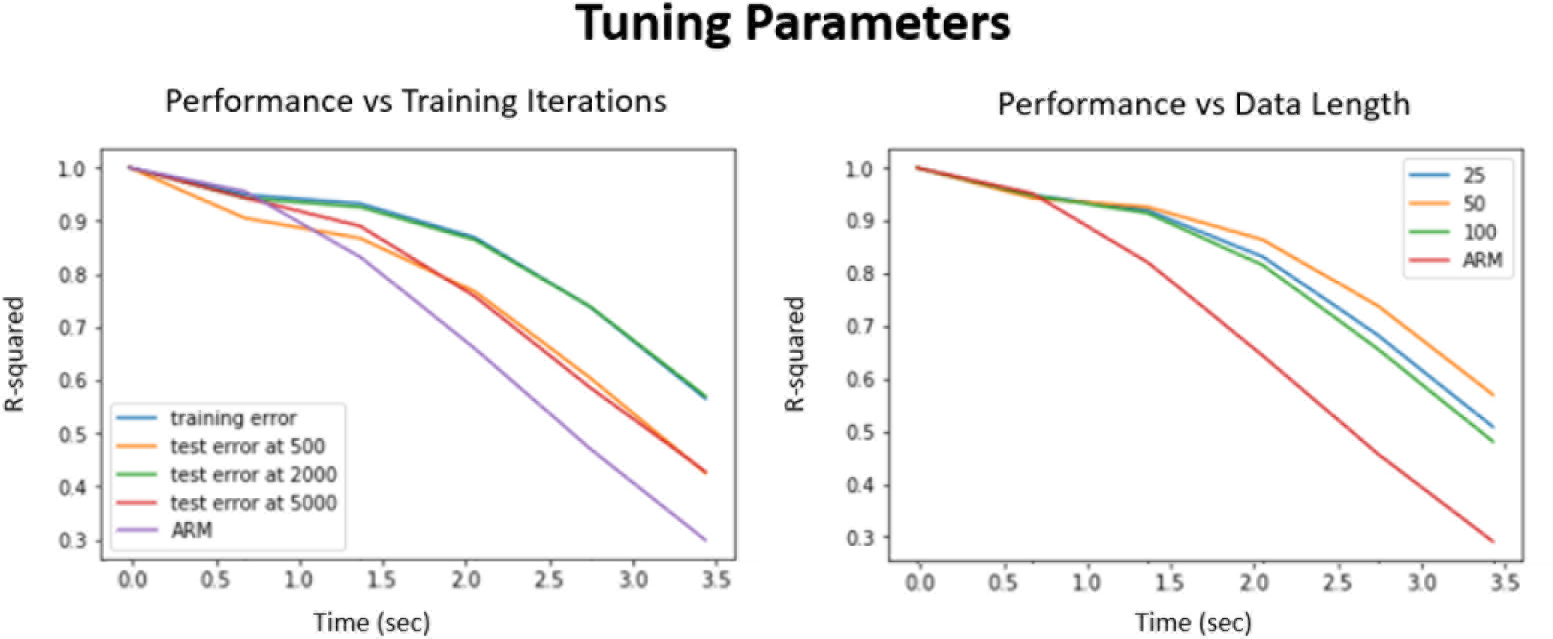
Tuning Parameters. Left: The effect of over and undertraining the network. The performance on the test data compared to the training data at 500 and 5000 is much worse. For our network size it performed best at around 2000 iterations. It is compared to the Autoregressive model (ARM) as baseline Right: The effect of picking different length segments and the performance accuracy. Again the maximum is closer to the middle which was in our case 50. Too small and too large networks either are slightly worse at learning the relationship between past and future rsfMIR timepoints

To speed up the training process, we utilized mini batches, where multiple instances of the training data are used simultaneously to train the network [19]. The number of instances that the network can be trained on simultaneously, depends on the size of the training data, and with 400 subjects we used 20 instances to simultaneously train the algorithm. The LSTM network in our model is initialized to a random point, and the first time segment supplies the initial state for the next segment. The performance on the very first block is very poor due to the unknown hidden state and is not included in our evaluation of the algorithm in the results section. This is a limitation with our implementation, and more complex architectures that solve for the initial state might circumvent this problem.

**Figure 8.**
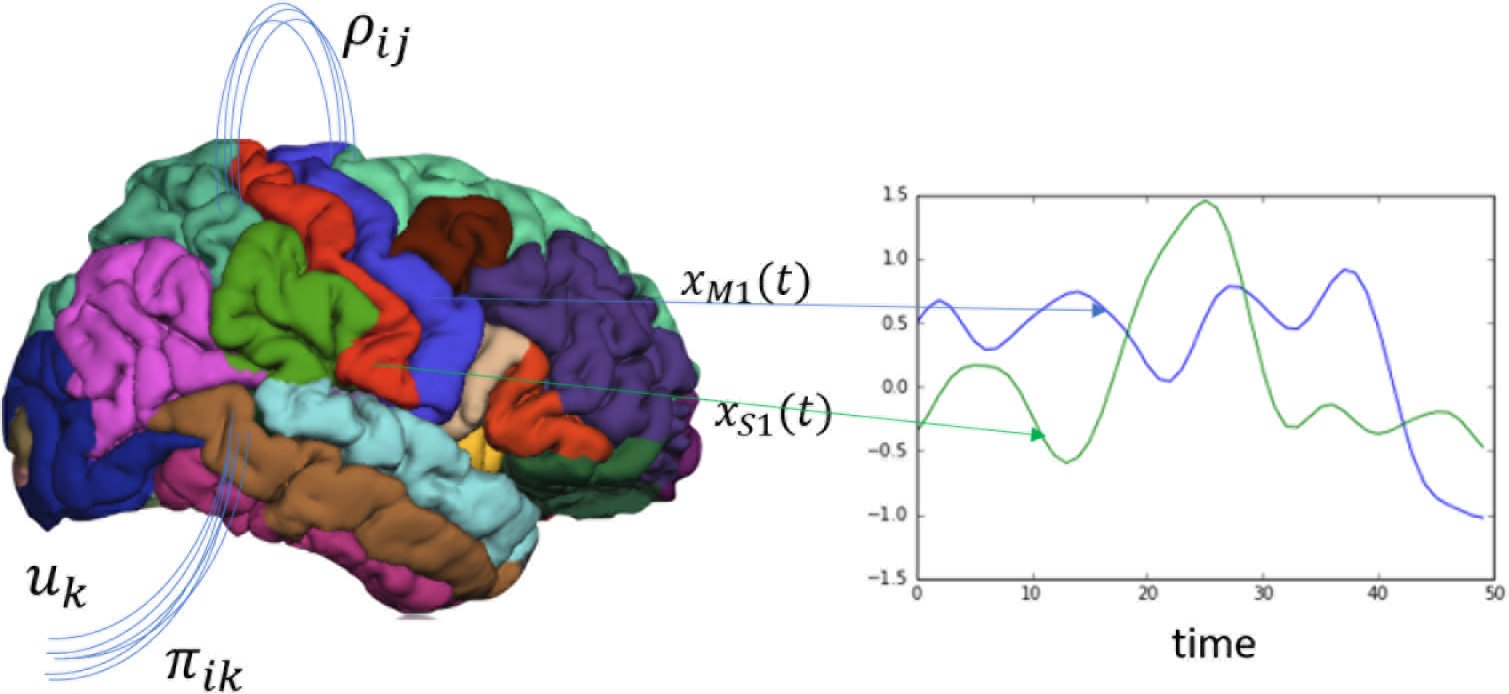
Brain Network Model. The Brain Network Model state space is constructed by averaging together the time courses of each parcellated region. The change of one of those areas *x*_*i*_ is a function of its own activity and its neighbors activity that it is connected with *ρ_ij_*, and the projection of external cortical input *u*_*k*_ to the brain via *π_ik_*.

### Brain Network Models

The Brain Network Model is constructed by specifying a parcellation or atlas, and each region of interest becomes the node and the edges represent the number of fibers between regions and is calculated using tractography. A Brain Network Model in its most general form, describes the change in neural activity x in region of interest i as a function of a sum of its neighbors j activity and its own activity and the physical properties of neural communication between i and j represented by the vector *ρ* (i.e the number of fibers between regions, the delay in propagation). The network dynamics are also mediated by a k-dimensional vector u representing all sub-cortical and sensory inputs, and the vector *π* representing again the physical properties that project these inputs into the brain (i.e thalamic tracts into cortex).

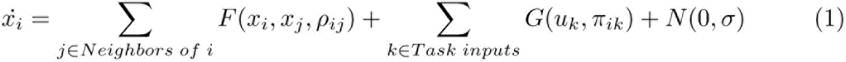

For resting state activity, the assumption is that *u*_*k*_(*t*) = 0 *t* and the first term dominates the activity. The function F for example can be as simple as the Firing Rate model

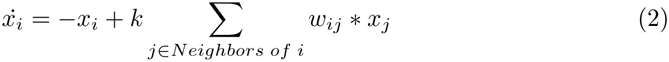

where *w*_*ij*_ represents the number of fibers between i and j, and k represents the global coupling parameter. In a more complex model the state variable x can also be represented by multiple variables such as the Wilson Cowan model shown in the equation below, which uses excitatory and inhibitory currents to describe the change in activity at every ROI. In the Firing Rate Model the output is taken to be the firing rate, and in the Wilson Cowan the fMRI signal is assumed to be just the excitatory signal since it dominates metabolically. These models are thus used to generate whole brain signals by choosing a random initial point and updating the next step via integration and generating the time series.

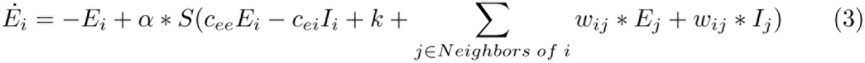

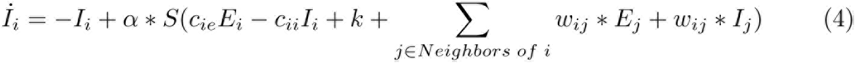

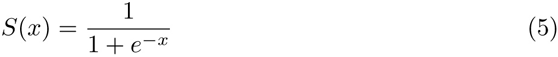

For our BNM, we choose the Firing Rate Model and the Wilson Cowan model described in more detail in Box 1. Our choice of Brain Network models reflects the constraints of this approach. Unlike the traditional BNM model where the simulation timestep can be arbitrary, we are constrained by the one-time step prediction approach of synchronizing with the measured data. Therefore, models that utilize high frequency oscillators such as the Hopf or the Kuramoto model and are usually simulated at a higher time resolution are more awkward to adapt into the fMRI framework, but would be more useful for training on faster datasets such as MEG or electroencephalogram (EEG). The two different models were also chosen in order to characterize the approach in terms of a simpler linear Firing Rate model and the more complex multi state nonlinear Wilson Cowan model. To account for noise in the BNM model, we chose to define the output of the LSTM as a distribution with a mean and standard deviation. We then sample from this distribution in order to generate the initial state. By representing the mapping as a non-deterministic process the algorithm generalizes to perform better on test data sets and gives more robust results between instantiations.

The output of the BNM is taken to be the next fMRI predicted timestep. The loss function then is taken as the difference between the predicted and the empirical next time points and the Autoencoder is trained based on this gradient. By forcing the output of the BNM to be the next predicted fMRI signal, the output of the LSTM is forced to become the closest initial time point and the LSTM solves for the non-linear transformation. We used the Tensor flow Adam Optimizer with a learning rate of 0.0001 to solve for the Autoencoder.

## Experimental Data

### Structural Network

To estimate the strucutral network we ran tractography on 5 HCP Diffusion Weighted Images using the freely available software Mrtrix [20, 34]. From the tractography we estimated the number of fibers that intersected two ROIs in the Desikan-Killiany atlas and normalized the power by dividing by the surface area of the receiving region [7]. The matrix is finally normalized by dividing by the largest eigenvalue in order for the graph Laplacian (k*SN-I) to have eigenvalues that are all negative [7]. This normalizes the dynamics so that the feedback decays over time, and does not exponentially increase the signal over time. The value of k is a hyperparameter, but simulations over a few different values around 0.9, showed that it made little difference, because the LSTM would just adjust its output correspondingly. The algorithm is robust as long as it is biased around values that would allow it to converge. For the Wilson Cowan we set both the k values to 0.9 as well, and learned the other parameters. We could also learn the value k, but since its not unique the reproduced latent state ends up further away from the signal. Since the autoencoder will fit the data either way, it is important to determine the constraints from the onset and constrain the latent state to be closer to the measurements.

### fMRI Data processing

Our resting state and task data is from the 447 minimally processed surface files from the Human Connectome project [34]. We took the MSMAII scans that were registered to standard space and in cifti format and ica-denoised them utilizing the 300 melodic ICA vectors that are provided from HCP. We transform from the surface-voxel time series to the ROI time series by averaging all voxels according to the parcellations established by the Desikan-Killiany atlas. This was done on an individual level since the surface parcellations are provided to by HCP and Freesurfer for each individual subject (aparc and aprac2009 files). The signal is then band passed filtered from 0.0008 Hz to 0.2 Hz and then global signal regressed using a general linear model with the mean time course of all cortical parcellations. The final signal is subsequently normalized along both axis [20]. For the task data, each dataset was processed separately (language, working memory, motor, social, emortional, gambling, relational) and then concatenated together. Each task dataset was rounded to the closest multiple of 50 and the autoencoder was fed alternating segments of task and the rest data. This signal is then fed as both the input and the output to the autoencoder and is the signal that we refer as the empirical rsfMRI for the rest of the paper [20].

### Dynamical analysis techniques

The BNA timeseries were first filtered (0.01 - 0.3Hz) before analyzing the properties using dynamical analysis techniques. The dynamical analysis techniques such as QPP and the k-Means analysis are described in detail in our previous publication, that outlines metrics in order to compare the simulated whole brain signal and the rsfMRI signals [20]. The QPP algorithm randomly picks a twenty second segment of data and correlates it with the whole signal. At the regions of peak correlation, the algorithm sums up all segments and creates a new template and iteratively converges to a repeating pattern [22]. The k-means analysis takes in sliding windowed (36 sec) functional connectivity matrices that are Fisher transformed and clusters them into seven different clusters [1]. We used an L1 distance to calculate the distance between matrices [1]. The resulting transitions between clusters was then quantified.

## Acknowledgements

This work has been supported by three grants. Part of this project was supported through a National Science Foundation Collaborative Research in Computational Neuroscience grant and the other part was supported through two National Institute Health RO1, N5078095 and a brain initiative grant MH111416.

We would like to thank the help of Dr. Chethan Pandarinath who provided invaluable help and insight with developing and interpretation of our initial Brain Network Autoencoder. We would also like to thank Dr. Christopher Rozell, for his insightful discussion on the interpretation of the Autoencoder.

